# SARS-CoV-2-neutralizing humoral IgA response occurs earlier but modest and diminishes faster compared to IgG response

**DOI:** 10.1101/2022.06.09.495422

**Authors:** Yuki Takamatsu, Kazumi Omata, Yosuke Shimizu, Noriko Kinoshita-Iwamoto, Mari Terada, Tetsuya Suzuki, Shinichiro Morioka, Yukari Uemura, Norio Ohmagari, Kenji Maeda, Hiroaki Mitsuya

## Abstract

Secretory immunoglobulin A (IgA) plays a crucial role in the mucosal immunity for preventing the invasion of the exogenous antigens, however, little has been understood about the neutralizing activity of serum IgA. Here, to examine the role of IgA antibodies against COVID-19 illnesses, we determined the neutralizing activity of serum/plasma IgG and IgA purified from previously SARS-CoV-2-infected and COVID-19 mRNA-vaccine-receiving individuals. We found that serum/plasma IgA possesses substantial but rather modest neutralizing activity against SARS-CoV-2 compared to IgG with no significant correlation with the disease severity. Neutralizing IgA and IgG antibodies achieved the greatest activity at approximately 25 and 35 days after symptom onset, respectively. However, neutralizing IgA activity quickly diminished and went down below the detection limit approximately 70 days after onset, while substantial IgG activity was observed till 200 days after onset. The total neutralizing activity in sera/plasmas of those with COVID-19 largely correlated with that in purified-IgG and purified-IgA and levels of anti-SARS-CoV-2-S1-binding IgG and anti-SARS-CoV-2-S1-binding IgA. In individuals who were previously infected with SARS-CoV-2 but had no detectable neutralizing IgA activity, a single dose of BNT162b2 or mRNA-1273 elicited potent serum/plasma neutralizing IgA activity but the second dose did not further strengthen the neutralization antibody response. The present data show that the systemic immune stimulation with natural infection and COVID-19 mRNA-vaccines elicit both SARS-CoV-2-specific neutralizing IgG and IgA response in serum, but the IgA response is modest and diminishes faster compared to IgG response.

**Author Summary:** Immunoglobulin A (IgA) is the most abundant type of antibody in the body mostly located on mucosal surfaces as a dimeric secretory IgA. Such secretory IgA plays an important role in preventing the adherence and invasions of foreign objects by its neutralizing activity, while monomeric serum IgA is thought to relate to the phagocytic immune system activation. Here, we report that individuals with the novel coronavirus disease (COVID-19) developed both systemic neutralizing IgG and IgA active against severe acute respiratory syndrome coronavirus 2 (SARS-CoV-2). Although the neutralizing IgA response was quick and reached the highest activity 25 days post-symptom-onset, compared to 35 days for IgG response, neutralizing IgA activity was modest and diminished faster than neutralizing IgG response. In individuals, who recovered from COVID-19 but had no detectable neutralizing IgA activity, a single dose of COVID-19 mRNA-vaccine elicited potent neutralizing IgA activity but the second dose did not further strengthen the antibody response. Our study provides novel insights into the role and the kinetics of serum IgA against the viral pathogen both in naturally-infected and COVID-19 mRNA-vaccine-receiving COVID-19-convalescent individuals.

## Introduction

Immunoglobulin A (IgA) is the most abundant type of antibody in the body [1], comprising most of the immunoglobulin in secretions primarily in the gut, milk, and bronchial secretions as a noninflammatory antibody against microbes [2]. Such secretory-IgA plays a crucial role in neutralizing the viruses, toxins, and inflammatory microbial molecules invading the mucosal epithelial cells [3] and exerts greater efficacy in preventing infections compared to serum IgG [4]. Thus, selective IgA deficiency, the most common immunologic defect in humans [5], causes recurrent sinopulmonary infections, autoimmune disorders, or allergic disorders. However, most individuals with selective IgA deficiency are asymptomatic and serum IgA levels in the patients do not necessarily correlate with the occurrence or severity of these disorders [6]. Serum IgA is the second most abundant isotype following IgG [7], and the functions of serum IgA appear to be related to the phagocytic system activation mediated through the Fc-alpha-RI (CD89) [8], although it has not been fully understood. In this regard, it had been recognized that the immunization via mucosal routes can elicit robust mucosal immune responses, while the systemic vaccination approach (*e*.*g*., administrated intramuscularly or intradermally) mainly induces IgG and apparently induces in part protective mucosal IgA responses [9].

In terms of the novel coronavirus disease (COVID-19), caused by severe acute respiratory syndrome coronavirus 2 (SARS-CoV-2), we previously reported that highly neutralizing activity-confirmed COVID-19 convalescent plasma and purified-IgG block the Syrian hamster disease progression with limited viral antigen-positive cells in terminal bronchioles and alveolar regions [10]. Sterlin *et al*. reported that mucosal IgA produced shortly after the symptom onset plays a crucial role in the early stage of the disease [11]. It has also been reported that COVID-19 mRNA-vaccines elicit high titer of anti-SARS-CoV-2-S1-binding IgG (S1-binding IgG) and IgA (S1-binding IgA) antibodies in serum [12-14].

In this regard, while systemic neutralizing IgG (nIgG) antibodies induced by COVID-19 and mRNA-vaccines are thought to be responsible for the protection against the symptomatic infection, further evaluation of the role of IgA in COVID-19 infection and COVID-19 vaccines, especially the evaluation of the neutralizing activity of such natural infection- or vaccine-induced IgA are needed.

Here, we report that individuals with COVID-19 developed both systemic nIgG and nIgA irrespective of the severity of the disease, however, even though the nIgA response was quick, the activity was modest and diminished faster compared to nIgG. We also report that the COVID-19 mRNA-vaccines elicit highly neutralizing serum IgA in COVID-19-experienced individuals.

## MATERIALS AND METHODS

### Participants

Fourteen individuals who were diagnosed with COVID-19 based on the positive RNA-quantitative-PCR (RNA-qPCR) results from February to April 2020 and eight individuals who received COVID-19 mRNA-vaccine (either BNT162b2 or mRNA-1273) from April to July 2021 after the recovery from COVID-19, and agreed to participate in the clinical studies (Certified Review Board of National Center for Global Health and Medicine approval numbers NCGM-G-003472 and NCGM-G-003536) for specimen collection and convalescent plasma donation [10,15] were enrolled in the present work. The data were analyzed anonymously. Nasopharyngeal swab samples were collected at early time points after admission and stored at -80°C until use. Sera or plasmas were obtained intermittently and stored at -20°C until use.

### Cells, viruses, and immunoglobulin purification

Transmembrane protease serine 2 (TMPRSS2)-overexpressing VeroE6 (VeroE6^TMPRSS2^) cells (RRID: CVCL_YQ49) were obtained from the Japanese Collection of Research Bioresources (JCRB) Cell Bank (Osaka, Japan). VeroE6^TMPRSS2^ cells were maintained in Dulbecco’s modified Eagle’s medium (DMEM) supplemented with 10% fetal bovine serum (FBS), 100 μg/ml penicillin, 100 μg/ml kanamycin, and 1 mg/ml G418 under a humidified atmosphere containing 5% CO_2_ at 37°C. A SARS-CoV-2 strain, SARS-CoV-2^05-2N^ (PANGO lineage B) was isolated in March 2020 in Tokyo, Japan as previously described [16]. IgG fractions were obtained from SARS-CoV-2-infected individuals’ sera or plasmas by using Spin column-based Antibody Purification Kit (Protein G) (Cosmo Bio, Tokyo, Japan). IgA fractions were purified from the IgG purification flow-through by using Pierce Jacalin Agarose (Thermo Fisher Scientific, Waltham, MA) and eluted in phosphate-buffered saline (PBS) by using Zeba™ Spin Desalting Columns, 40K MWCO (Thermo Fisher Scientific). The total human IgG and IgA concentrations were determined by using the Human IgG ELISA Kit and Human IgA ELISA Kit, respectively (abcam, Cambridge, UK). The purity of the IgG and IgA was determined by using the capillary electrophoresis Simple Western Jess apparatus and the Total Protein Detection Module (Protein Simple, San Jose, CA), Anti-Human IgA, alpha-Chain Specific, HRP-Linked Antibody #80403, and Anti-Human IgG, Fc gamma Fragment Specific, HRP-Linked Antibody #32935 (Cell Signaling Technology, Danvers, MA). The purities of the IgG and IgA were approximately 85% (84.0 ± 2.4) and 75% (75.2 ± 1.6), respectively as four representative IgG and IgA samples were examined.

### Antiviral assays

The SARS-CoV-2 neutralizing activity of donated plasma and purified immunoglobulin against the wild-type SARS-CoV-2 (PANGO lineage B) was determined as previously described [10,16,17]. In brief, VeroE6^TMPRSS2^ cells were seeded in 96-well flat microtiter culture plates at the density of 1 × 10^4^ cells/well. On the following day, the virus (SARS-CoV-2^05-2N^) was mixed with the various concentrations of the serum/plasma or purified immunoglobulin fractions and incubated for 20 min. at 37°C. The preincubated mixture was inoculated to the cells at a multiplicity of infection (MOI) of 0.01. The cells were cultured for 3 days and the number of viable cells in each well was measured using Cell Counting Kit-8 (Dojindo, Kumamoto, Japan). The potency of SARS-CoV-2 inhibition by sera/plasmas or purified immunoglobulin was determined based on its inhibitory effect on virally-induced cytopathicity in VeroE6^TMPRSS2^ cells. The amounts of S1-binding antibodies in each plasma sample were determined by using Anti-SARS-CoV-2 ELISA (IgG) and (IgA) (Euroimmun, Lübeck, Germany). The serial diluted donor 84 (D84) plasma [10] was used as a reference (100%) for quantification with four parameters logistic curve calculated by using Image J (Fiji) (**S1 Fig**.) [18].

### Statistical analysis

The 50% neutralizing titers of sera/plasmas (NT_50_), 50% effective concentration of purified-IgG and -IgA (nIgG-EC_50_ and nIgA-EC_50_, respectively), and the amounts of anti-SARS-CoV-2-S1-binding-IgG and anti-SARS-CoV-2-S1-binding-IgA (S1-binding IgG and S1-binding IgA, respectively) were determined and compared between the acute and convalescent phases of COVID-19 and between the moderate and severe symptoms using Wilcoxon signed-rank test and Wilcoxon rank sum test, respectively. The attenuation rates of nIgG-EC_50_ and nIgA-EC_50_ were calculated by dividing the nIgG-EC_50_ or nIgA-EC_50_ values determined the latest in the study with the highest neutralizing activity (lowest nIgG-EC_50_ or nIgA-EC_50_ values) by days 28, 42, and 56 post-onset. To examine which of nIgG-EC_50_ and nIgA-EC_50_ values diminished faster in the convalescent-vaccine group, the values obtained by subtracting the lowest EC_50_ values from the highest EC_50_ values post-1^st^ vaccine administration were compared. Then, the attenuation rates of nIgG-EC_50_ and nIgA-EC_50_, the slopes made with the first and second S1-binding IgA and IgG amounts, and the differences after the vaccination were compared by Wilcoxon signed-rank test. The correlations and corresponding 95% confidence intervals of NT_50_, IgG-EC_50_, IgA-EC_50_, S1-binding IgG, and S1-binding IgA were determined using the repeated measures correlation method to consider the within-individual association [19] using rmcorr R package ver. 0.4.6 [20]. The computed correlation coefficients were considered high if the absolute value was above 0.7, moderate if the absolute value was between 0.4 to 0.7, and low if the absolute value was below 0.4, according to Guilford’s Rule of Thumb. The nIgG-EC_50_ and nIgA-EC_50_ kinetics were fitted with a Generalized Additive Model [21] with mgcv R package ver.1.8-40. The fitting was implemented for superimposed data of all samples. All the analyses were performed using R statistical software ver. 4.1.3 [22]. Statistical significance was defined as *p*< 0.05.

## Results

### Clinical characteristics of the participants

Fourteen individuals, who were confirmed to have SARS-CoV-2 infection with positive RNA-quantitative-PCR (RNA-qPCR) results and admitted to the Center Hospital of the National Center for Global Health and Medicine in Tokyo, Japan from February to April 2020 (COVID-19 group) (**Table 1**), and eight individuals, who received COVID-19 mRNA-vaccine (either BNT162b2 or mRNA-1273) from April to July 2021 after the recovery from COVID-19 (convalescent-vaccine group) (**Table 2**), were enrolled. These individuals agreed to participate in the present clinical studies. All the individuals were Japanese and 2 out of 14 (14.3%) in the COVID-19 group and 2 out of 8 (25.0%) in the convalescent-vaccine group were female (**Tables 1, 2**). The median (range) age was 53 (37 to 68) and 53 (35 to 61) years in the COVID-19 group and convalescent-vaccine group, respectively (**Tables 1, 2**). In the COVID-19 group, seven individuals (50%) had moderate symptoms of lower respiratory disease or imaging with no oxygen requirement, while seven individuals (50%) had severe symptoms and required oxygen treatment during the clinical course without any sequential organ failure. There were no significant differences in the age, sex, or sample collection dates between the moderate and severe symptom groups (**Table 1**). Individuals in the COVID-19 group received experimental therapeutic agents, which are now mostly considered to be ineffective (**S1 Table**) [23]. The convalescent-vaccine group received the primary series of COVID-19 mRNA-vaccine 70 to 458 (median 306) days after the disease onset (**Table 2**).

**Table 1.** The characteristics of COVID-19 experienced individuals (COVID-19 group) in the study.

|  | All patients<br>(n = 14) | Moderate<br>(n = 7) | Severe<br>(n = 7) |
| --- | --- | --- | --- |
| Age, years (median) | 37 – 68 (53) | 47 – 62 (53) | 37 – 68 (63) |
| Sex |  |  |  |
| Male | 12 (85.7%) | 6 (85.7%) | 6 (85.7%) |
| Female | 2 (14.3%) | 1 (14.3%) | 1 (14.3%) |
| Oxygen Requirement |  |  |  |
| day (median) |  | N/A | 2 – 15 (6.5) |
| None | 7 (50.0%) | 7 (100%) | 0 (0%) |
| Nasal Canula | 6 (42.9%) | 0 (0%) | 6 (85.7%) |
| High-Flow Nasal Canula | 1 (7.1%) | 0 (0%) | 1 (14.3%) |
| Sample collection, day (median) |  |  |  |
| Acute phase | 3 – 15 (9) | 3 – 13 (8) | 4 – 15 (9) |
| Convalescent phase | 17 – 201 (86) | 19 – 201 (88) | 17 – 196 (74) |

**Table 2.** The characteristics of COVID-19 mRNA-vaccinee after the recovery from the disease (convalescent-vaccine group) in the study.

|  | All participants<br>(n = 8) | mRNA-1273<br>(n = 3) | BNT162b2<br>(n = 5) |
| --- | --- | --- | --- |
| Age, years (median) | 35 – 61 (53) | 55 – 61 (55) | 35 – 56 (44) |
| Sex |  |  |  |
| Male | 6 (75.0%) | 2 (66.6%) | 4 (80.0%) |
| Female | 2 (25.0%) | 1 (33.3%) | 1 (20.0%) |
| Severity |  |  |  |
| Mild | 5 (62.5%) | 2 (66.6%) | 3 (60.0%) |
| Moderate | 2 (25.0%) | 0 (0%) | 2 (40.0%) |
| Severe | 1 (12.5%) | 1 (33.3%) | 0 (0%) |
| Days to vaccination after onset (median) | 70 – 458 (306) | 173 – 458 (436) | 70 – 335 (286) |
| Sample collection, day (median) |  |  |  |
| Before 1st vaccination (median) | 17 – 298 (209) | 97 – 249 (209) | 17 – 298 (219) |
| After 1st vaccination (median) | 4 – 103 (56) | 4 – 91 (57) | 8 – 103 (55) |

#### SARS-CoV-2-neutralizing sera/plasmas IgA response occurs earlier and diminishes faster compared to IgG response

We previously described the kinetics of neutralizing activity of immunoglobulin G (IgG) fractions purified from plasmas of 43 SARS-CoV-2-infected individuals using cell-based assays [16]. In the current study, we chose fourteen individuals with moderate to severe COVID-19 symptoms and evaluated neutralizing activity of whole sera/plasmas and purified-IgG and -IgA fractions against wild-type SARS-CoV-2^05-2N^ (PANGO lineage B). As shown in Fig 1A, whole sera/plasmas from all fourteen SARS-CoV-2-infected individuals had significantly high titers of SARS-CoV-2-neutralizing activity by 30 days after symptom onset, and thereafter their neutralizing activity gradually decreased but the decay became slower after around 50 days (**Fig 1A**). Significant levels of neutralizing activity persisted in sera/plasmas in all participants as examined on up to day 200 (**Fig 1A**). Purified-IgG from sera/plasmas also exerted neutralizing activity expressed as 50% effective concentration (EC_50_) of up to 1.0 μg/mL (**Fig 1B**) and showed substantial neutralizing activity by around 200 days post-onset. Substantial amounts of S1-binding IgG antibodies were also seen by around 200 days after the onset (**Fig 1E**). We also identified good immune response to produce nIgA following the emergence of COVID-19 symptoms, however, the decay of the IgA neutralizing activity occurred much earlier than that of nIgG, and by 129 days after the onset, 12 of the 14 individuals (85.7%) had around EC_50_ value of 100 µg/mL or undetectable (>100 µg/mL) neutralizing activity (**Fig 1C**). Three of the 14 individuals (21.4%) showed no detectable nIgA activity throughout the study. In contrast to the early decay in nIgA activity, substantial amounts of S1-binding IgA persisted by up to 200 days after the onset (**Fig 1F**).

**Fig 1.**
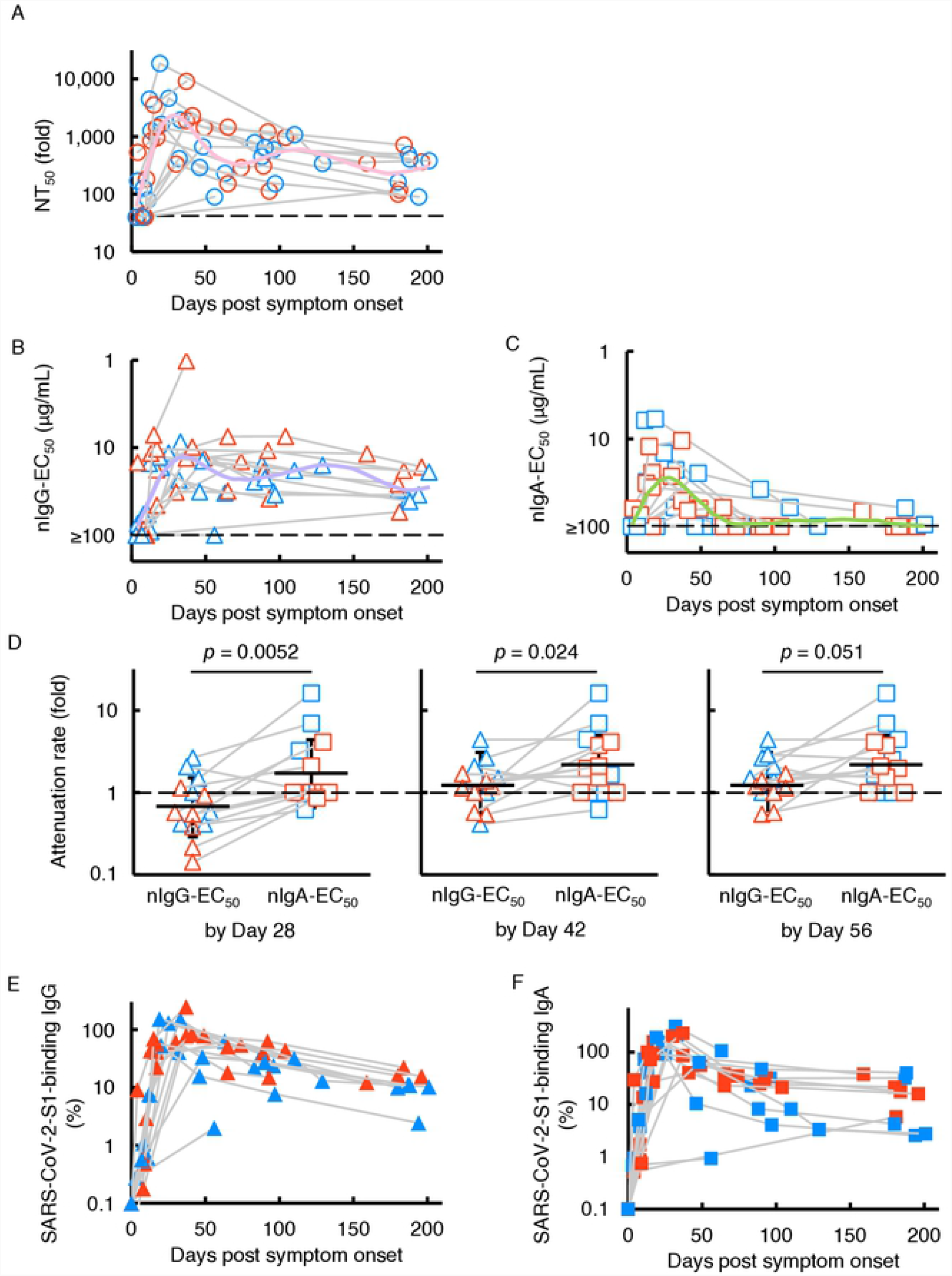
Kinetics of SARS-CoV-2-neutralizing activity and S1-binding antibodies. VeroE6^TMPRSS2^ cells were exposed to wild-type SARS-CoV-2^05-2N^ with or without various concentrations of diluted sera/plasmas (**A**), purified-IgG (**B**), or purified-IgA (**C**), and the neutralizing activity and the amounts of S1-binding antibodies were determined. The dashed line denotes the assay limit values (≤40-fold for the panel a and ≥100 μg/mL for panels b and c). Note that the highest viral neutralizing activity of purified-IgG and -IgA was seen around 35 and 25 days after onset, respectively. Furthermore, the neutralizing activity of serum IgA diminished much quicker than that of IgG. The colored line (NT_50_ for pink, nIgG-EC_50_ for purple, and nIgA-EC_50_ for light green) denote the fitted curve. (**D**) Attenuation rate of the nIgG-EC_50_ and nIgA-EC_50_ between the highest neutralizing activity of purified IgG and IgA by day 28, 42, and 56 post-onset and neutralizing activity determined the latest in the study. The kinetics of the amount of S1-binding IgG (**E**) and IgA (**F**) were also shown. The amount of S1-binding IgG and IgA increased approximately by day 21 post symptom onset, followed by a gradual decrease. Note that by contrast, substantial amounts of S1-binding-IgG and - IgA persisted around 200 days after the onset, while the decay occurred more rapidly in IgA.

To quantify and compare the time-dependent kinetics of sera/plasmas, nIgG, and nIgA activity, we generated fitted curves by using Generalized Additive Model [21,24], which showed that nIgA response occurred significantly earlier than nIgG response; it took 25 days post-onset for nIgA response to reach its peak but 35 days for nIgG response to reach its peak (**Fig 1B** and **C**). It was also noted that the nIgA response diminished faster than nIgG response; the average nIgA-EC_50_ value virtually reached the detection limit (≥100 μg/mL) approximately 70 days post-onset (**Fig 1C**), while substantial nIgG activity persisted until ∼200 days post-onset **(Fig 1B)**. We also attempted to quantify the time-dependent reduction of nIgG-EC_50_ and nIgA-EC_50_ by calculating the attenuation rate between the highest neutralizing activity of purified-IgG and -IgA by day 28 (range 3-25), 42 (range 3-41), and 56 (range 3-56) post-onset and neutralizing activity determined the latest in the study (**Fig 1D**). As shown in Fig 1D, the attenuation rates of nIgA-EC_50_ were significantly greater than those of nIgG-EC_50_ by 28 (4 weeks; **Fig 1D**, left panel) and 42 (6 weeks; **Fig 1D**, middle panel) days post-onset with *p* values of 0.0052 and 0.024, respectively. The same trend was seen when the attenuation rates of nIgG-EC_50_ and nIgA-EC_50_ were determined by 56 days (8 weeks; **Fig 1D**, right panel, *p*=0.051).

Sterlin and his colleagues have previously reported that IgA dominates the early neutralizing antibody response to SARS-CoV-2 in patients with COVID-19 [11]. Thus, we attempted to examine whether the S1-binding IgA production predominated timewise over the S1-binding IgG production by using the slopes made with the first and second S1-binding IgA and IgG amounts determined in each individual. The comparative data showed that S1-binding IgA production significantly predominated over S1-binding IgG production (*p*=0.009, Wilcoxon signed-rank test) (**Fig 1E** and **1F**).

### Neutralizing activity is greater in patients with severe COVID-19 than with moderate disease

We next asked if higher neutralization activity is seen in acute (less than 14 days post-onset or when oxygen treatment was required) or convalescent (14 days post-onset and when no oxygen was required) phase or in patients with moderate or severe COVID-19. A significant increase was seen in 50% neutralizing titers (NT_50_) of sera/plasmas in convalescent phase than in acute phase in both moderate and severe symptom groups (*p*=0.02 and 0.03, respectively) (**Fig 2A**). There was also a significant increase in nIgG activity in convalescent phase in both moderate and severe symptom groups (*p*=0.03 and 0.03, respectively) (**Fig 2B**). The same pattern was seen in S1-binding IgG amounts (**Fig 2D**). By contrast, there was no significant difference in nIgA activity between acute or convalescent phases in either moderate or severe symptom groups (**Fig 2C**). The amounts of S1-binding IgA were higher in convalescent than in acute phase in moderate symptom group, although the difference in the severe symptom group was not significant (**Fig 2E**).

**Fig 2.**
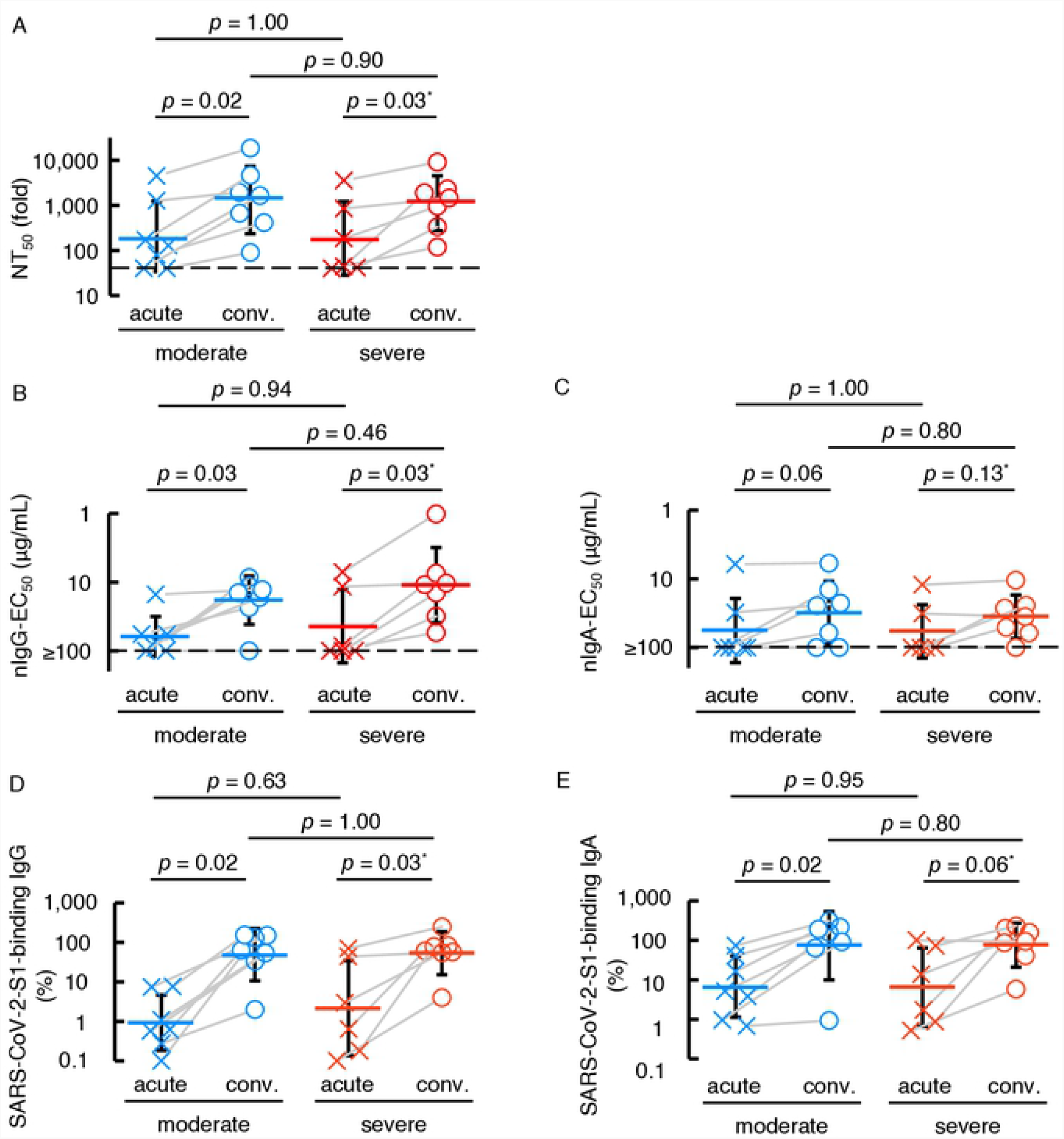
COVID-19-convalescent individuals possess the greater neutralizing activity and SARS-CoV-2-S1-binding antibody levels than those at the acute phase. The neutralizing activity of sera/plasmas, purified-IgG, and purified-IgA (**A, B**, and **C**, respectively) and the amounts of S1-binding IgG and IgA (**D** and **E**, respectively) were compared between the acute phase (less than 14 days post-symptom onset or when the individual required oxygen treatment) and the convalescent phase (14 days post-symptom onset and beyond with no oxygen requirement).

### Contribution of neutralizing IgG antibody to sera/plasmas neutralizing activity is greater than that of neutralizing IgA

The amounts of S1-binding IgG antibodies in sera from patients with COVID-19 highly correlate with SARS-CoV-2-specific neutralizing activity levels in serum IgG fraction [16,25]. However, the role of SARS-CoV-2-specific humoral IgA antibodies in protecting against SARS-CoV-2 infection remains clarified. Thus, we asked whether S1-binding IgA antibody amounts correlate with nIgA activity. The NT_50_ values of sera/plasmas from patients with COVID-19 proved to well correlate with nIgG activity (nIgG-EC_50_ values) with the rho (ρ) value of -0.72 (95% confidence interval [CI]; -0.84 to -0.54) (**Fig 3A**), in line with our previous observations [16]. In the case of purified-IgA from patients with COVID-19, high correlation was also observed with nIgA activity (nIgA-EC_50_ values) with the ρ value of -0.78 (95% CI; -0.88 to -0.62), although 31 of 56 IgA samples had very low or undetectable (≥100 µg/mL) neutralization activity (**Fig 3B**). Between nIgG-EC_50_ and nIgA-EC_50_ values, however, there was a moderate correlation was seen with the ρ value of 0.42 (95% CI; 0.13 to 0.65) (**Fig 3C**). The NT_50_ values of sera/plasmas and nIgG-EC_50_ values also had high correlation with S1-binding IgG amounts (**S2A** and **S2B Fig**). There was also high correlation between the NT_50_ values of sera/plasmas and nIgA-EC_50_ values with S1-binding IgA amounts (**S2C** and **S2D Fig**). S1-binding IgA in nasopharyngeal swab samples collected at the earliest point of the infection (less than 20 days post symptom onset) tend to have a higher amount as the day goes (**S3A Fig**). Further, the nasal S1-binding IgA was highly correlated with serum S1-binding IgA with Spearman’s ρ value of 0.73 (95% CI; 0.40 to 0.89) (**S3B Fig**). On the other hand, the serum total human IgG and IgA were consistent during the study period (**S2C** and **S2D Fig**) with a low correlation (**S2E Fig**).

**Fig 3.**
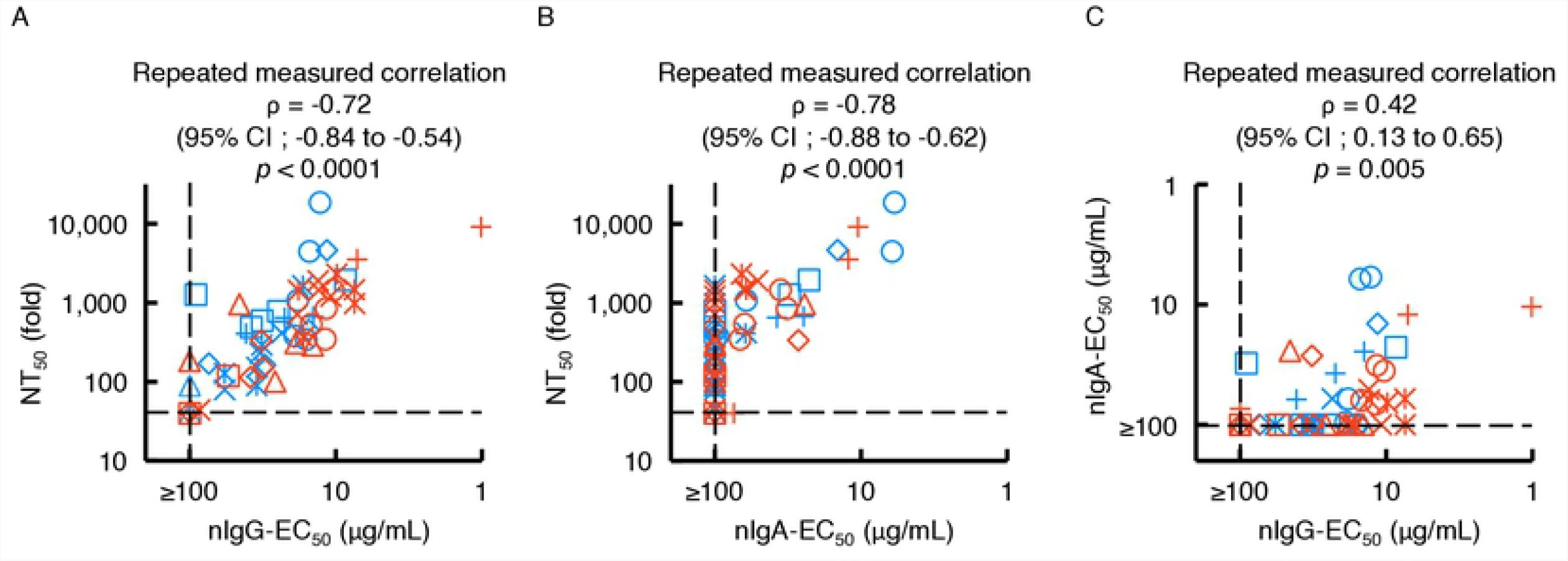
Correlations of purified-IgG and -IgA neutralizing activities with sera/plasmas neutralizing titers. The neutralizing activity of purified-IgG (**A**, nIgG-EC_50_) and -IgA (**B**, nIgA-EC_50_) against sera/plasmas neutralizing activity (NT_50_) values are plotted. (**C**) The nIgA-EC_50_ values are plotted against the nIgG-EC_50_ values. Note that a high correlation is observed between NT_50_ values and nIgG-EC_50_ values (Repeated measured correlation ρ = -0.72 (95%CI; -0.84 to - 0.54) (**A**) and between the NT_50_ values and NIgA-EC50 values (ρ = -0.78 (95%CI; -0.88 to - 0.62) (**B**), while moderate correlation was observed between nIgA-EC_50_ and nIgG-EC_50_ (Repeated measured correlation ρ = 0.42 (95%CI; 0.13 to 0.65) (**C**). Each symbol denotes the sample from one and the same individual.

The present data suggest that the neutralizing activity seen in sera/plasmas of patients with COVID-19 is largely composed of the neutralizing activity of serum IgG (**Fig 3A**) but also of that of IgA (**Fig 3B**). Moreover, as has been seen in the case of neutralizing activity of sera/plasmas that is in large correlated with the amount of S1-binding IgG [16,20], the neutralizing activity of serum IgA is correlated with the amounts of S1-binding IgA (**S2D Fig**), while the neutralizing activity of IgA was modest compared to that of IgG (**Fig 1B, 1C**, and **3C**).

### mRNA-COVID-19 vaccine induces high-level neutralizing activity in COVID-19 convalescent individuals

We next examined the SARS-CoV-2-specific IgG and IgA neutralizing activity elicited with the primary series of mRNA vaccine administration (BNT162b2 or mRNA-1273) in eight individuals who had experienced qPCR-confirmed symptomatic COVID-19 70 to 458 days before the first immunization (**Table 2**). All eight individuals had low but detectable to moderate levels of neutralizing activity in sera/plasmas before the vaccination (**Fig 4A**). Most of these individuals had significantly high titers of neutralizing activity within 28 days after the first vaccination. The high NT_50_ titers were not further boosted following the second dose, which is a quite different pattern of NT_50_ values from the patterns seen in those who were SARS-CoV-2-naïve and received the first and second doses of vaccine [17,26,27]. A similar pattern was seen when nIgG-EC_50_ values were determined in the same participants (**Fig 4B**). In the case of nIgA-EC_50_ values, none of the participants had detectable neutralizing activity (≥100 µg/mL) before the COVID-19 mRNA-vaccination, but 3 of the 8 participants had a substantial rise in the nIgA-EC_50_ values before the second dose of mRNA-vaccine (**Fig 4C**). In contrast, these COVID-19-experienced individuals had moderate to high levels of S1-binding IgG and IgA before the first dose, and greater levels of S1-binding IgG and IgA were documented following the first dose although no further increase was seen after the second dose (**Fig 4D** and **4E**). It was noted however that the nIgA activity rapidly decreased (**Fig 4C**) compared to nIgG activity (**Fig 4B**), although such rapid decay was not seen in the amounts of S1-binding IgA antibodies (**Fig 4E**).

**Fig 4.**
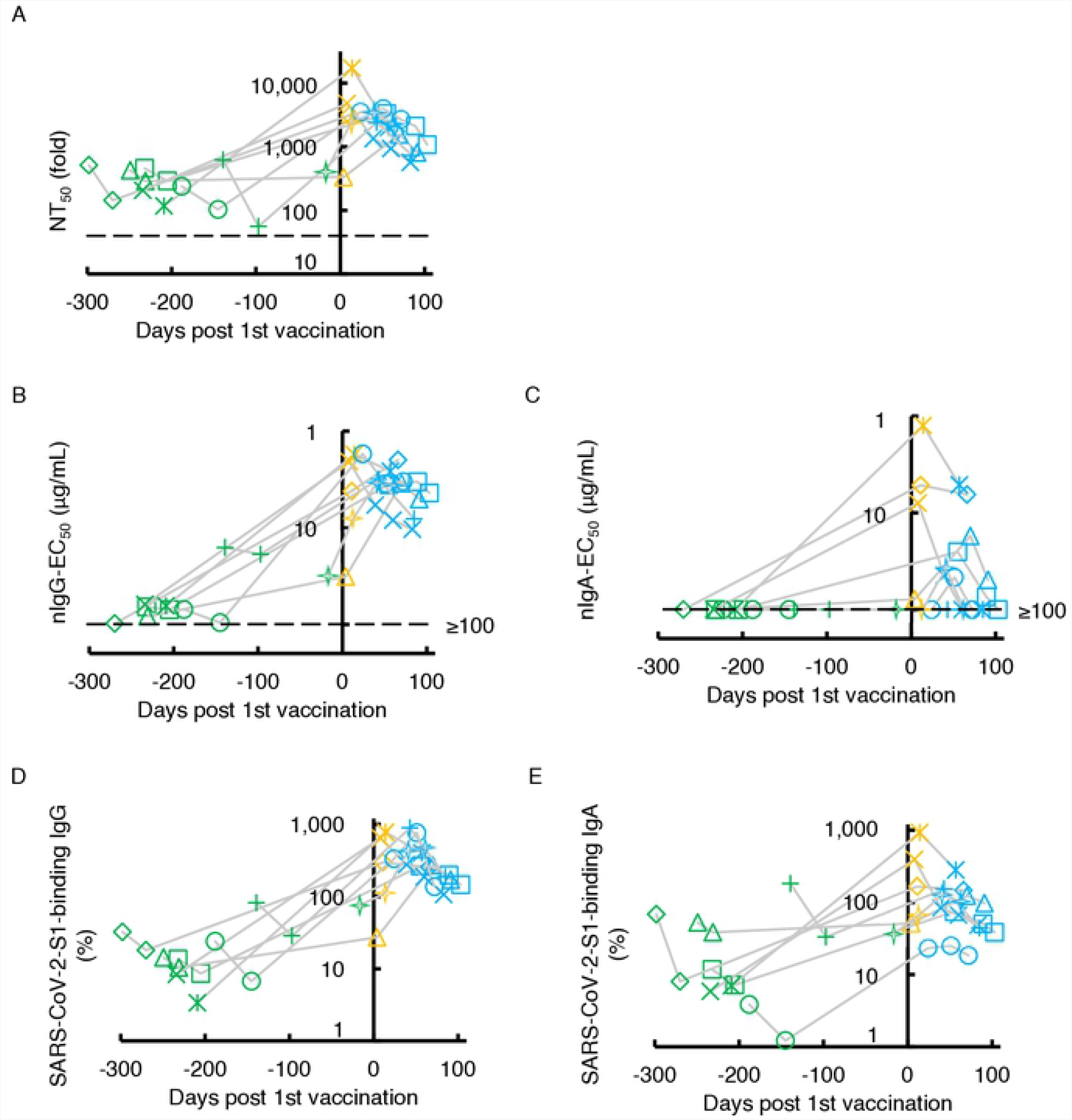
Kinetics of neutralizing activity and S1-binding antibody levels before and after COVID-19 mRNA-vaccination. The kinetics of neutralizing activity (**A, B**, and **C**) and S1-binding antibody levels (**D** and **E**) in eight previously COVID-experienced individuals who received the COVID-19 mRNA-vaccine are shown. Note that all the values significantly rose after the first dose of the vaccine. Also note that none of the participants had detectable nIgA-EC_50_ values (≥100 µg/mL) before vaccination (**C**). On the other hand, all of them had low to high levels of nIgA-EC_50_ after a singled dose of the vaccine and such levels quickly decreased (**C**) and their S1-binding IgA levels persisted after the two doses of vaccine in the study period (**E**). The dashed line denotes the assay detection limit (≤40-fold dilution for NT_50_ and ≥100 μg/mL for nIgG-EC_50_ and nIgA-EC_50_). Green symbols denote the samples collected before COVID-19 mRNA-vaccination, while yellow and light-blue denote after the 1^st^ and 2^nd^ doses, respectively. Each symbol denotes the sample from one and the same individual.

All of the NT_50_, nIgG-EC_50_, nIgA-EC_50_, % S1-binding IgG, and % S1-binding IgA values proved to have significantly increased following primary series administration (**S4A-E Fig**). These data demonstrate that COVID-19 mRNA-vaccines induce high titers of nIgA in previously COVID-19-experienced individuals after a single dose of vaccine. However, such nIgA activity apparently diminished faster compared to nIgG activity, while the difference was not statistically significant (*p*=0.069) (**Fig 4B** and **4C**). Such an early decay of nIgA activity had been seen in those with symptomatic infection with SARS-CoV-2 (**Fig 1C**). The second dose vaccination did not significantly slow the speed of decay (**Fig 4C**). We also examined whether there are correlations among NT_50_, nIgG-EC_50_, nIgA-EC_50_, and S1-binding IgG and IgA values. As we have seen that nIgG activity greatly contributes to sera/plasmas SARS-CoV-2-neutralizing activity compared to that of nIgA activity in individuals with COVID-19 (**Fig 3A-C** and **S2A-D Fig**), similar profiles were identified in previously-COVID-19-contracted individuals following COVID-19 mRNA vaccination (**S5A-G Fig**).

## DISCUSSION

In respiratory tract infections such as influenza virus infection, natural infection induces systemic IgG responses [28,29] as well as mucosal secretory-IgA responses [30]. In terms of COVID-19, Sterlin *et al*. reported that IgA-expressing circulating plasmablasts were detected shortly after the symptom onset which have a consistent phenotype with that found in lung and eventually produced mucosal IgA [11]. Our present data also showed that the amounts of nasal S1-binding IgA antibody and the amounts of serum S1-bindng IgA are highly correlated (Pearson’s ρ=0.73, **S3B Fig**), suggesting the serum S1-binding IgA and mucosal S1-binding IgA share similar, albeit not the same, antigenic determinants or immunological repertoire in response to SARS-CoV-2-S1. In this regard, it is of note that Wang *et al*. have suggested that serum IgA monomers are produced by the same cells that produce secretory dimers [25]. Of note, Sterlin *et al*. reported that serum IgA specific to the receptor-binding domain (RBD), which represents a critical target for neutralization, was detected earlier than anti-RBD IgG as assessed with a photonic ring immunoassay [11]. In the present study, we also showed that nIgA response occurred significantly earlier than nIgG response; it took 25 days post-onset for nIgA response to reach its peak and 35 days for nIgG response to reach its peak (**Fig 1B** and **C**). Moreover, S1-binding IgA production significantly predominated over S1-binding IgG production (**Fig 1D** and **E**). These data are in line with the observations by Sterlin *et al*. [11]

Although the neutralization of pathogens is attributed to the neutralizing activity of IgG, providing long-term immunity for as long as decades, such as mumps, varicella-zoster virus (VZV), and Epstein–Barr virus (EBV) [31], protective immunity to seasonal coronaviruses [32], SARS-CoV, and Middle East respiratory syndrome (MERS)-CoV [33] is known to be short-lived. Vanshylla *et al*. reported that the neutralizing activity in serum waned quickly (half-life; 3.6 months) compared to the neutralizing activity of purified-IgG (half-life; 7.8 months), and such a short half-life of activity of serum is thought to be partially attributed to the presence of S-binding IgA and IgM in serum [34]. Moreover, Iyer *et al*. reported RBD-binding IgA antibodies are short-lived compared to RBD-binding IgG antibodies [35]. In the present study, we extended the observations by Vanshylla *et al*. and Iver *et al*. and demonstrated that SARS-CoV-2-neutralizing activity of sera/plasmas IgA was identified earlier and diminished faster than that of IgG as assessed in 14 individuals with COVID-19 (**Fig 1B** and **C**). Moreover, when such activity was determined following mRNA vaccination in eight COVID-19-experienced individuals, SARS-CoV-2-neutralizing activity in sera/plasmas IgA also quickly diminished as compared to that in sera/plasmas IgG (**Fig 4B** and **C**). However, there were no significant differences in the decay rate of SARS-CoV-2-S1 binding IgG and IgA levels (**Fig 1D, 1E, 4D**, and **4E**). The faster decay in the nIgA activity compared to that in nIgG may derive from the difference in half-lives of serum IgA and IgG (*i*.*e*., 3-5 and 21 days, respectively). Also, it is possible that since the total amount of serum IgG in the body is greater than that of IgA, the consumption and absorption of neutralizing IgA by the viral antigens could be more apparent than in the case of IgG.

It has been reported that the neutralizing activity of IgM and IgA are dramatically greater than that of IgG when the activity of recombinant monoclonal antibodies, which share the same anti-SARS-CoV-2-spike protein Fab region, was examined using a pseudo-typed lentivirus coated with the SARS-CoV-2 spike protein and angiotensin converting enzyme 2 (ACE2)-transfected Crandell-Rees feline kidney cells as the host cell line [36]. In the current study, unlike their findings, we observed that the neutralizing activity of purified-IgA is modest compared to that of purified-IgG (**Fig 1B, 1C**, and **3C**). In this regard, we have used a cell-based neutralization assay using IgA fractions purified from sera/plasmas, which are of polyclonal nature. Thus, our data should possibly represent the more comprehensive protective effect of serum-derived IgA, although more studies are needed.

There are reports that individuals with selective IgA deficiency tend to have higher risks of severe COVID-19 [37,38]. Thus, we initially hypothesized that individuals with moderate symptoms would possess greater neutralizing activity of serum IgA than those with severe COVID-19. However, there were no significant differences in nIgA-EC_50_ values between those with moderate and severe diseases (**Fig 2D** and **2E**). In this regard, we have lately shown that patients with severe COVID-19 had greater nIgG levels in serum than those with mild COVID-19 [16] and we reasoned that the exposure to larger amounts of SARS-CoV-2 over long-term in those with severe COVID-19 resulted in the greater nIgG activity [34].

It should be noted that the limitation in the present work is that we did not systematically characterize SARS-CoV-2-specific secretory IgA antibodies, which represent the dominating immunoglobulins in exocrine secretions. In the literature, there are currently only a few reports documenting the role of secretory IgA antibodies in protection against SARS-CoV-2 infection. Studies to elucidate the protective effect of secretory IgA upon SARS-CoV-2 infection and anti-COVID-19 vaccination remain to be conducted.

In conclusion, the present data showed that SARS-CoV-2-neutralizing serum/plasma IgA response is seen earlier than nIgG response, suggesting that the humoral IgA plays a critical role in the acute phase of the infection, although that nIgA response diminishes faster compared to nIgG response, which should in turn play a role in the later phase of infection. Further, in previously SARS-CoV-2-infected individuals, the first (initial) administration of COVID-19 mRNA-vaccines induces high titers of nIgG as well as nIgA, however, the neutralizing activity of IgA also diminishes faster than that of IgG.

## Contributors

Conceptualization, Y.T. and H.M.; Methodology, Y.T., K.M., and H.M.; Formal Analysis, K.O., Y.S., and Y.U.; Investigation, Y.T., K.O., and N.K-I.; Data curation, Y.T., Y.S., N.K-I., M.T., and T.S.; Resources, N.K-I., M.T., T.S., and S.M.; Writing – Original Draft, Y.T. and H.M.; Writing-Review & Editing, K.O., Y.S., N.K-I., Y.U., and K.M.; Supervision, N.O., K.M., and H.M.; Project Administration, H.M.; Funding Acquisition, K.M. and H.M.

## Acknowledgments

This work was supported in part by Japan Agency for Medical Research and Development (AMED) (grant numbers JP20fk0108160 and JP20fk0108502 to K.M. and JP20fk0108502, JP20fk0108257, and JP20fk0108510 to H.M.); by MHLW Research on Emerging and Re-emerging Infectious Diseases and Immunization Program (grant number JPMH20HA1006 to K.M.); by a grant from National Center for Global Health and Medicine Research Institute (grant number 20A2003D to K.M.); and by the Intramural Research Program of the Center for Cancer Research, National Cancer Institute, National Institutes of Health (H.M.). The funders had no role in study design, data collection and analysis, decision to publish, or preparation of the manuscript. We thank Mariko Kato for technical assistance. We would like to thank all the patients who participated in the clinical trial for the collection of convalescent plasma and staff of the Diseases Control and Prevention Center, Center for Clinical Sciences, Department of Hematology, Clinical Laboratory Department, Department of Clinical Engineering, and Nursing Department at the Central Hospital of National Center for Global Health and Medicine.

## Supporting Information

**S1 Fig. Four parameters curve fit model of the quantification of S1-binding antibody levels using the commercially available S1-binding IgA ELISA**.

**S2 Fig. High correlations of purified-IgG and -IgA neutralizing activities with S1-binding antibody levels**.

The NT_50_ values against S1-binding IgG and IgA levels are shown in panels **A** and **C**, respectively, and nIgG-EC_50_ and nIgA-EC_50_ values against the S1-binding IgG and IgA are shown in panels **B** and **D**, respectively.

**S3 Fig. Kinetics and the correlations of nasal SARS-CoV-2-S1-binding-IgA levels and total IgG and IgA amounts in serum**.

The % SARS-CoV-2-S1-binding IgA levels in nasal swab samples were determined with the commercially available S1-binding IgA ELISA using a COVID-19-convalescent plasma’s S1-binding IgA that was referred as 100%. (**A**) Temporal changes of the nasal S1-binding-IgA levels in over 18 days following the onset of the disease. (**B**) Correlation of % nasal S1-binding-IgA levels with that of sera/plasmas S1-binding IgA. Temporal changes of total human IgG and IgA levels following the diseases (**C** and **D**). Correlation of total human IgA levels with that of IgG is shown (**E**).

**S4 Fig. COVID-19 mRNA-vaccine induces significant neutralizing activity and S1-binding antibody levels in COVID-19-experienced individuals**.

The neutralizing activity of sera/plasmas, purified-IgG, and purified-IgA (**A, B**, and **D**, respectively) and the amounts of S1-binding IgG and S1-binding IgA (**C** and **E**, respectively) were compared between the pre- and post-vaccination.

**S5 Fig. Correlations of sera/plasmas, purified-IgG, and -IgA neutralizing activities with S1-binding antibody levels**.

The NT_50_ values against (**A**) nIgG-EC_50_ values, (**B**) nIgA-EC_50_ values, (**D**) S1-binding-IgG level (S1-binding IgG), and (**F**) S1-binding-IgA level are plotted. Note that neutralizing activity of IgG primarily contributes to sera/plasmas SARS-CoV-2-neutralizing activity compared to that of IgA (**A, B**, and **C**) in previously-COVID-19-contracted individuals following COVID-19 mRNA vaccination.

**S1 Table.** Experimental therapeutic agents used in the COVID-19 group.

|  | All patients<br>(n = 14) | Moderate<br>(n = 7) | Severe<br>(n = 7) |
| --- | --- | --- | --- |
| Experimental therapeutic agents |  |  |  |
| Remdesivir (RDV) | 4 (28.6%) | 2 (28.6%) | 2 (28.6%) |
| Lopinavir/ritonavir (LPV/r) | 2 (14.3%) | 1 (14.3%) | 1 (14.3%) |
| Hydroxychloroquine (HCQ) | 3 (21.4%) | 2 (28.6%) | 1 (14.3%) |
| HCQ + Azithromycin (AZM) | 3 (21.4%) | 0 (0%) | 3 (42.9%) |
| inhaled Ciclesonide (CIC) | 1 (7.1%) | 1 (14.3%) | 0 (0%) |
| Favipiravir (FPV) | 1 (7.1%) | 0 (0%) | 1 (14.3%) |
| None | 2 (14.3%) | 1 (14.3%) | 1 (14.3%) |
| Corticosteroid use |  |  |  |
| Hydrocortisone (HDC) | 4 (28.6%) | 0 (0%) | 4 (57.1%) |
| Methylprednisolone (mPSL) | 1 (7.1%) | 0 (0%) | 1 (14.3%) |
| PMX-DHP | 3 (21.4%) | 0 (0%) | 3 (42.9%) |

Abbreviation: PMX-DHP; polymyxin B-immobilized fiber column direct hemoperfusion

